# Utility of T1 mapping and T2 mapping for non-invasive assessment of liver fibrosis: preclinical results

**DOI:** 10.1101/2023.11.08.566354

**Authors:** Jing Rong, Yujie Zhu, Kun Zhu, Min Shao, Xiujuan Yin, Tongtong Liu, Xiao Wang

**Author notes:** **Corresponding author:** GN: Xiao FN: Wang, PhD, Department of Radiology, the First Affiliated Hospital of Anhui Medical University, Hefei, Anhui 230022, China. Jingwen Luo and Zhiwei Zhang contributed equally to this work (Jing Rong, Yujie Zhu).

## Abstract

**Objective:** To explore the potential of T1 and T2 mappings in assessing liver fibrosis (LF) and investigate the relationships between MRI and liver fibrogenesis markers.

Materials and methods A total of 39 male C57BL/6 mice were divided into the control group (n = 9) and the model group (n = 10 per subgroup) with carbon tetrachloride (CCl4) administration for 2, 4, and 6 weeks. The METAVIR system (F0-4) was performed to stage fibrosis. MRI T1 and T2 mappings were performed and T1, and T2 values were calculated. One-way analysis of variance (ANOVA), Spearman’s rank correlation analysis, and receiver operating characteristic (ROC) curves were performed.

**Results:** T1 and T2 values increased with progressing severity of fibrosis induction (*P* < 0.01). T1 and T2 were significantly correlated with LF stages (ρ = 0.854, 0.697, *P* < 0.001). The area under the curves (AUCs) range of T1 and T2 for predicting ≥F1, ≥F2, ≥F3, and F4 were 0.842-0.994 and 0.808-0.883, respectively. T1 and T2 showed moderate to strong correlations with collagen-associated protein and inflammatory factors.

**Discussion:** T1 and T2 mappings can evaluate and differentiate LF stages in the CCl4-induced model. T1 is better correlated with collagen deposition and inflammation of LF than T2.

## Introduction

Liver fibrosis (LF) is regarded as a healing response caused by excessive deposition of collagen-based extracellular matrix (ECM) following repeated liver injury. Depending on etiologies, LF is associated with several fundamental pathological processes, such as activation of hepatic stellate cells (HSCs), collagen accumulation, inflammation reaction, etc[1]. HSC activation and their trans-differentiation to myofibroblasts (MFB) is a major hallmark of LF[2]. With the progression of LF, the expression of related liver fibrogenesis markers will significantly upregulate[3]. Although the exact mechanism of LF is poorly understood in many cases, it is considered a dynamic process with the potential for partial or complete regression[4]. Therefore, an accurate evaluation of early-stage LF is imperative to prevent and even reverse the process of fibrosis.

To date, liver biopsy remains the gold standard for diagnosing and staging LF[5]. However, it presents several limitations such as high intra/interobserver variability, risk of sampling error, invasive and poorly reproducible operation with potential adverse complications[6]. Liver-related serum markers are widely used in the clinic because of their noninvasive and reproducible nature, but most of them have low sensitivity and specificity[7]. Consequently, reliable and noninvasive imaging techniques are urgently required in clinical studies.

Recently, noninvasive imaging techniques for quantifying and diagnosing LF were reported, including MR elastography, Diffusion-weighted imaging, R2* mapping, Extracellular volume mapping, etc. all of which are of particular importance[8–11]. Relaxation times (eg, T1, T2) are inherent parameters of tissues and were reported to provide high diagnostic accuracy in the assessment of LF[12, 13]. However, widespread clinical adoption of T1 and T2 mappings has been limited by a lack of consensus findings in the assessment of LF[14–16]. Furthermore, the corresponding liver fibrogenesis markers of collagen accumulation and inflammation reaction, such as hydroxyproline (HYP), α-smooth muscle actin (α-SMA), interleukin-1 (IL-1), interleukin-6 (IL-6), and tumor necrosis factor-α (TNF-α), may be closely related to T1 and T2[11, 17]. However, as far as we know, there have been few studies well characterized the relationship between T1, and T2 with these liver fibrogenesis markers.

The purpose of the present study was thus to explore the potential of T1 and T2 mappings in assessing LF and investigate the relationships between MRI parameters and liver fibrogenesis markers in a CCl4-induced mice model.

## Materials and methods

### Liver fibrosis model in mice

A total of 39 male C57BL/6 mice (ca. 20–25g, 6 weeks old) were used for this study and randomly divided into the model (M, n = 30) and control groups (N, n = 9). The LF model was induced by intraperitoneal injection of 10% CCl4 olive oil mixed suspension twice a week at a dose of 0.05 ml/10g for 2, 4, and 6 weeks. To characterize the histopathologic changes, extracellular protein expression of LF, and imaging of the model, the model groups after injection for 2, 4, and 6 weeks were designated as three different time point subgroups (n = 10 per subgroup). The mice in the control group received injections of saline solution at the same pattern, dosage, and frequency (n = 3 per time point). All mice were randomly selected at the time points of 2, 4, and 6 weeks for MRI examination.

### MR imaging acquisition

Examinations were performed using a 1.5T MRI scanner (Ingenia, Philips Healthcare, Best, The Netherlands) with an 8-channel mouse coil. The mice were anesthetized by intraperitoneal injection of 0.5mg/10g sodium pentobarbital solution. The chest and abdomen were wrapped with a manual bandage to reduce the influence of respiratory motion artifacts. For the determination of T1 and T2 values, T1 and T2 mappings were acquired. Detailed descriptions of MRI sequences and parameters were provided in Electronic Supplementary Material Table S1.

### Imaging analysis

Quantitative T1 and T2 maps were automatically reconstructed using the image-processing workstation (InterlliSpace™ Portal, Philips) after data acquisition. T1 maps were computed by applying the equations described by Messroghli et al, where three-parameter nonlinear curve fitting using a Levenberg-Marquardt algorithm was performed on a pixel-by-pixel basis[18]. T2 maps were computed by the mono-exponential fitting of the multi-echoes TSE signals using a least-square nonlinear algorithm on a pixel-by-pixel basis[19]. For each mouse, the layer comprised of the largest homogeneous liver region where the middle hepatic vein merged into the inferior vena cava was selected, and three regions of interest (ROIs) measuring 8.0 mm² were placed in the right, middle and left lobes of the liver, respectively. ROIs were performed by two radiologists (with 15-year and 4-year experience in abdomen imaging work, respectively) who were blinded to the histopathological results. The hepatic blood vessels, biliary tract, liver edge, and obvious artificial artifacts were avoided, and the average T1, and T2 values were recorded and used for final analysis.

### Histopathological analysis

All the mice were sacrificed for postmortem verification after MRI scanning. Liver samples were fixed in 10% formaldehyde for 24 hours and embedded in paraffin. Liver slices were stained with hematoxylin and eosin (HE), Masson collagen (Masson), and Sirius red to evaluate LF. Two experienced pathologists assessed all liver specimens, who were blinded to the MRI results. Any confused histological degrees were resolved by consensus. LF staging (F0-4) was assessed according to the Meta-analysis of Histological Data in Viral Hepatitis for staging LF (METAVIR) classification system[20]: F0 = no fibrosis; F1 = mild fibrosis, portal area fibrosis without septum formation; F2 = substantial fibrosis, periportal fibrosis with a little fiber septum formation; F3 = advanced fibrosis, massive septal formation without cirrhosis; F4 = widespread fibrosis, early cirrhosis. Briefly, early-stage fibrosis started in F1 and extended in F2 from hepatic central venules (C). Stage F3 belonged to the middle or late phases of LF, characterized by a large number of fibrous septa formed and lobular structures but no pseudolobules. Stage F4 showed that C-C septa connected and formed into pseudolobules.

### Liver fibrogenesis markers quantification

The amounts of α-SMA, IL-1, IL-6, and TNF-α protein in mice were measured by Western Blot as previously described[21]. The primary antibodies were specific for the following proteins: α-SMA (Bioss, Beijing, China); IL-1, IL-6 and TNF-α (WANLEIBIO, Shenyang, China). The Protein expression levels were normalized with β-actin (internal control) expression, and fold changes in expression were calculated. Hepatic HYP level was measured photometrically in liver hydrolysates according to the instructions of the hydroxyproline content detection kit (Solarbio, Beijing, China). Results are expressed in μg/g liver.

### Statistical analysis

Data were reported as mean ± SD (standard deviation). Statistical analysis was performed using IBM SPSS Statistics (v.25.0, Chicago, IL, USA), GraphPadPrism (v.9.3.0, San Diego, California), and MedCalc (v.20.0, Mariakerke, Belgium). By calculating the intra-class correlation coefficient (ICC) to assess the repeatability of measurements, and ICC ≥ 0.75 was considered a good consistency. If the data were normally distributed with homogeneity of variance, one-way ANOVA followed by the least significant difference (LSD) test was performed. Otherwise, the Kruskal–Wallis test followed by the Nemenyi test was performed. Receiver operating characteristic (ROC) curve analysis was used to assess the diagnostic performance of MRI parameters for staging LF. The area under the curve (AUC), sensitivity, specificity, and cut-off value were calculated, and the AUC values were compared between T1 and T2 using DeLong’s test. The correlation between quantitative MRI parameters, fibrosis stages, and liver fibrogenesis markers was assessed using Spearman’s rank correlation analysis. Correlation coefficients were stratified as follows: poor = 0-0.19; fair = 0.20-0.49; moderate = 0.50-0.69; very strong = 0.70-0.89; and perfect = 0.90-1.00[22]. *P* < 0.05 was indicated as statistically significant.

## Results

### Repeatability

The ICCs of T1 and T2 values measured by the two readers were 0.980 (*P* < 0.001, 95% confidence interval: 0.935-0.994) and 0.869 (*P* < 0.001, 95% confidence interval: 0.627-0.958), which showed T1 shared similar good repeatability with T2 values.

### Histopathological analysis

The survival rate was 92.3% (36/39) in the model group and all mice in the control group survived. Compared with normal liver specimens, the liver of the 6-week model group appeared granular, indicating that the LF model was successfully prepared (Fig 1a, b). Histological analysis demonstrated that fibrosis development in the model group maintained an upward trend with significance (Fig 1c). Typical H&E, Masson, and Sirius red staining for stage F0-4 after CCl4 administration was shown in Fig 1d. In the model groups, the fibrosis stages for the 27 mice were as follows: F1 = 5, F2 = 7, F3 =7, and F4 = 8. In the control group, the fibrosis stages of all 9 mice were F0.

**Figure 1.**
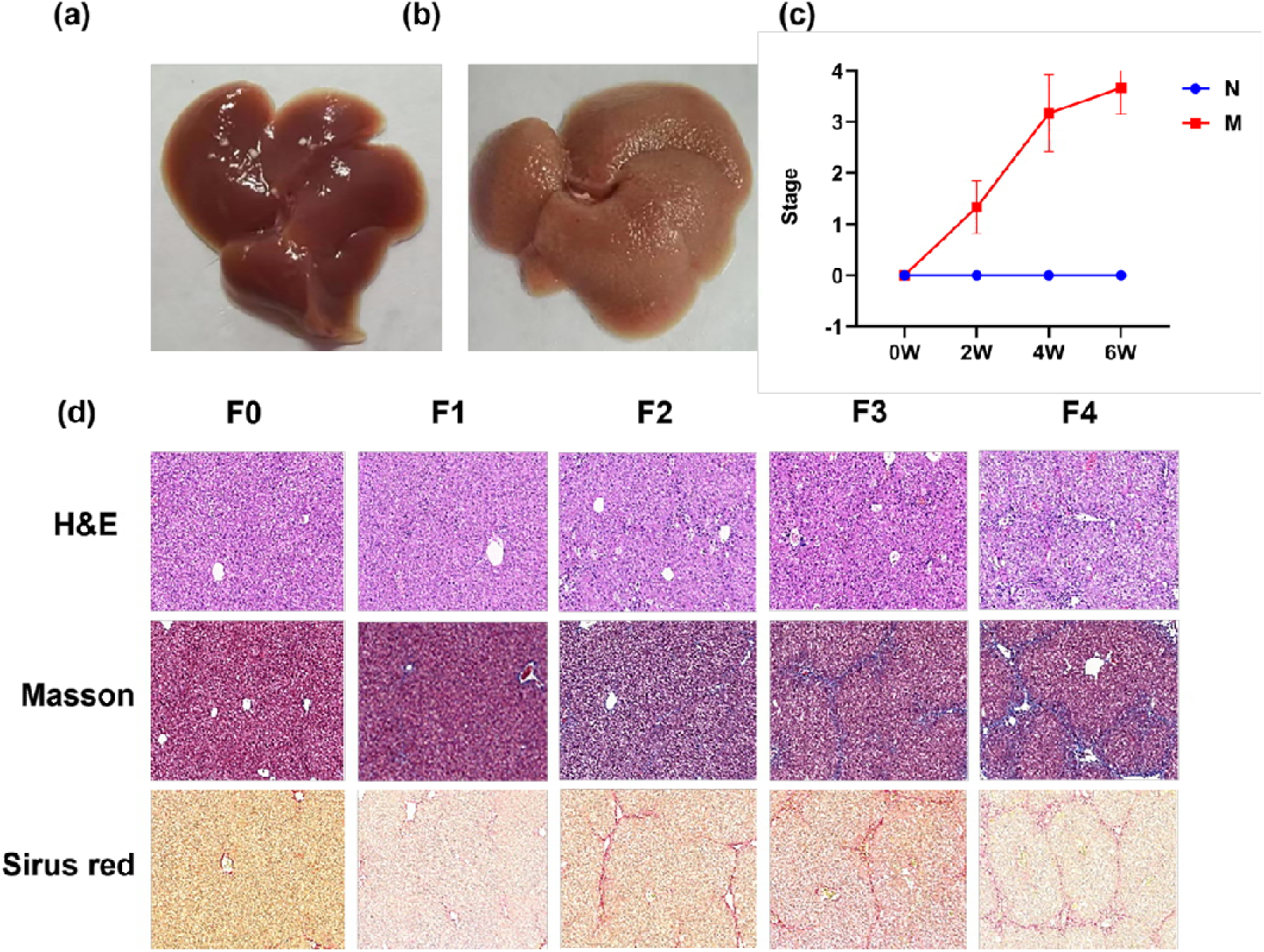
Images of model and pathological analysis. (a) The liver specimen of mice in the control group. (b) The liver specimen of the 6-week model group. (c) Distribution of fibrosis stages in control (N) and model (M) groups at different time points. LF development in the M group maintained an upward trend with significance (ρ=0.947, *P* < 0.001). (d) Representative images of H&E, Masson, and Sirius red staining of the liver tissue for METAVIR F0-4 stages. (light microscopy, ×200 original magnification). H&E, hematoxylin and eosin.

### T1, T2, and liver fibrogenesis markers at different timepoints

As shown in Table 1, T1 values increased with progressing severity of fibrosis induction in the model subgroups at 2, 4, and 6 weeks, and significant differences between fibrosis subgroups as a whole were observed (*P* < 0.01), although there was an overlap between groups. Similar trends were observed in the model subgroups for the T2 values (*P* < 0.01), and a significant difference in T2 values was found between week 2 and week 6 (*P* = 0.024). As shown in Table 1 and Electronic Supplementary Material Fig. S1, a continuous increase in liver fibrogenesis markers (HYP, α-SMA, IL-1, IL-6, and TNF-α) was observed, although the differences of α-SMA and IL-1 between week 2 and week 4, week 4 and week 6 were not statistically significant (all *P* > 0.05).

**Table 1:** General characteristics of the different subgroups of mice.

| Variable | Control (n<br>= 9) | 2 Weeks<br>(n =10) | 4 Weeks<br>(n = 10) | 6 Weeks<br>(n = 10) | P-Value |
| --- | --- | --- | --- | --- | --- |
| T1 values (ms) | 498.12 ± 36<br>.27 | 552.92 ± 34.<br>16<sup>\$</sup> | 645.27 ± 32.2<br>6<sup>\$#</sup> | 682.68 ± 37.39<br>\$# | < 0.00<br>1 |
| T2 values (ms) | 38.19 ± 2.5<br>8 | 42.13 ± 1.88 | 44.15 ± 5.20<sup>\$</sup> | 47.86 ± 6.03<sup>\$#</sup> | 0.002 |
| **Collagen-associated protein** |  |  |  |  |  |
| HYP content (μ<br>g/g) | 194.44±7.5<br>3 | 352.65±9.35<sup>\$</sup> | 420.14±2.08<sup>\$#</sup> | 549.06±12.83<sup>\$</sup><br>#& | < 0.001 |
| α-SMA (fold change) | 1.05±0.05 | 1.58±0.10<sup>\$</sup> | 1.67±0.08<sup>\$</sup> | 1.75±0.09<sup>\$#</sup> | < 0.001 |
| **Inflammatory factors** |  |  |  |  |  |
| IL-1 (fold change) | 1.08±0.08 | 1.48±0.10<sup>\$</sup> | 1.62±0.16<sup>\$</sup> | 1.88±0.23<sup>\$</sup> | 0.001 |
| IL-6 (fold change) | 1.03±0.04 | 1.24±0.02<sup>\$</sup> | 1.40±0.09<sup>\$#</sup> | 1.57±0.05<sup>\$#&</sup> | < 0.001 |
| TNF-α (fold change) | 1.09±0.09 | 1.37±0.03<sup>\$</sup> | 1.58±0.04<sup>\$#</sup> | 1.69±0.06<sup>\$#&</sup> | < 0.001 |

*P*-Value represents the comparison between the model and control groups.
<sup>\$</sup>*P* < 0.05 vs. Control; <sup>#</sup>*P* < 0.05 vs. 2 weeks; <sup>&</sup>*P* < 0.05 vs. 4 weeks.
HYP, hydroxyproline; α-SMA, α-smooth muscle actin;
IL-1, interleukin-1; IL-6, interleukin-6; TNF-α , tumor necrosis factor-α .

### T1 and T2 in the assessment of fibrosis stages

As shown in Fig 2, Fig 3, and Electronic Supplementary Material Table S2, T1 values tended to increase with the extent of LF as a whole (*P* < 0.001) and demonstrated a significant positive correlation with the LF stages (ρ = 0.854, *P* < 0.001). Pairwise comparisons showed that the differences in T1 values between F0-2 vs. F3-4 were statistically significant (all *P* < 0.01). Similar trends were observed for T2 values with increasing degrees of fibrosis in general (*P* = 0.003), and there was a moderate positive correlation (ρ = 0.697, *P* < 0.001) between T2 values and LF stages. Furthermore, ROC analysis was conducted to determine the potential of T1, and T2 values (Table 2). The AUC ranges of T1, and T2 for diagnosing ≥F1, ≥F2, ≥F3, and F4 were 0.842-0.994, and 0.808-0.883 respectively. No significant difference in diagnostic performance was found between T1 and T2 values (all *P* > 0.05).

**Figure 2.**
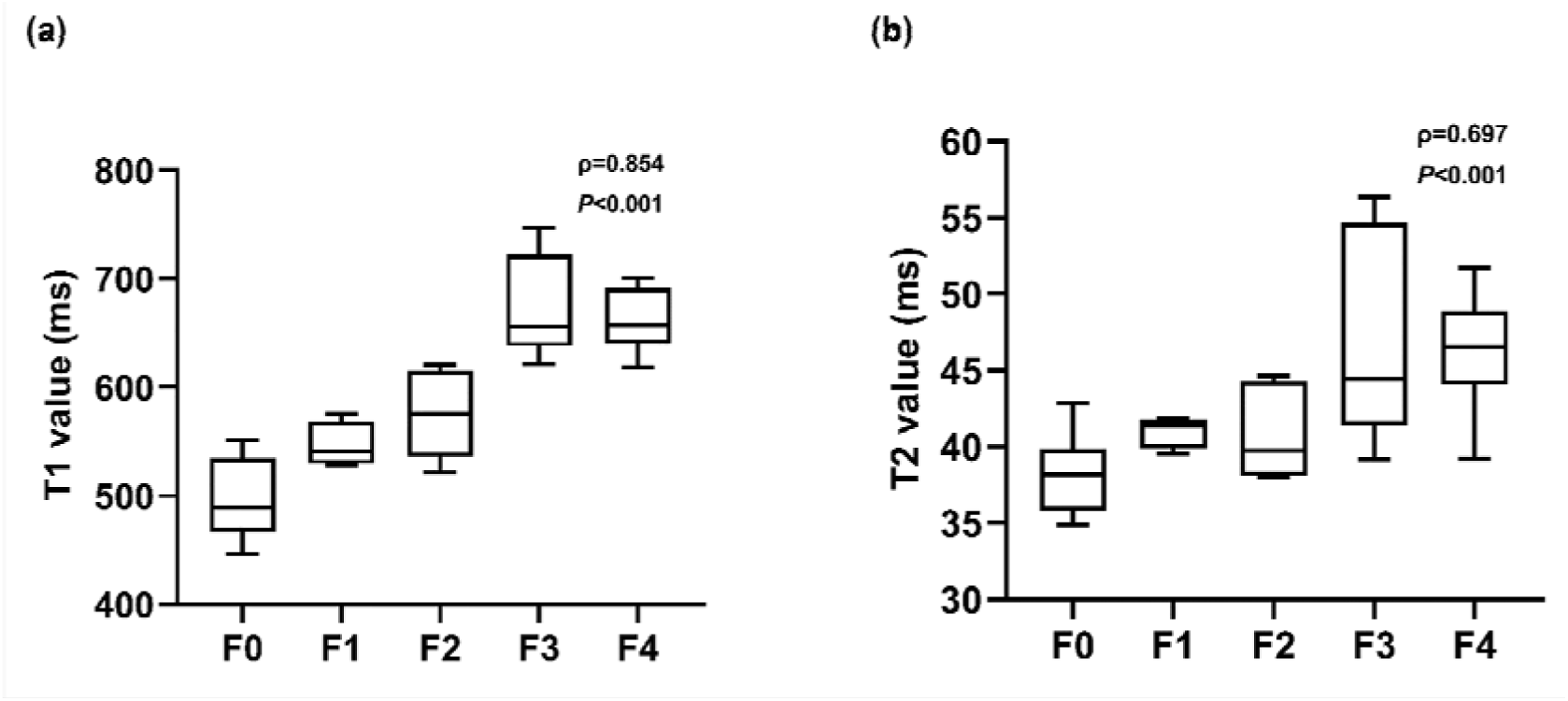
Box charts of T1 (a) and T2 (b) values in different fibrosis stages, and Spearman’s rank correlation between MRI parameters and fibrosis stages.

**Figure 3.**
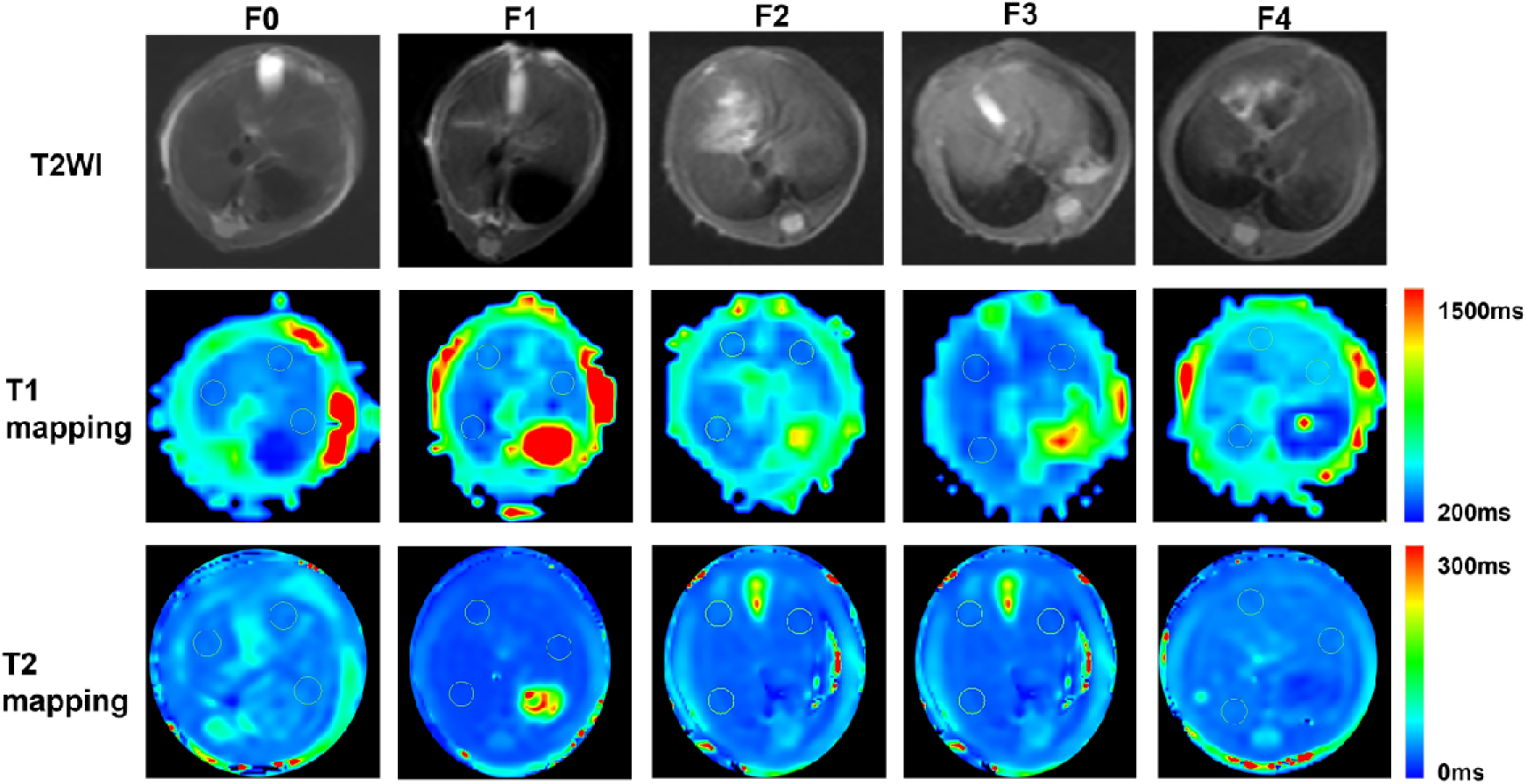
Representative axial images of T2WI, T1 maps and T2 maps in the control (F0) and model (F1-4) groups. Three ROIs measuring 8.0 mm² are located on the largest slice of the liver area of MR images. The average T1 values of F0-4 maps are 489.32ms, 533.08ms, 575.89ms, 654.25ms, 688.45ms, respectively. The average T2 values of F0-4 maps are 35.22ms, 39.54ms, 44.02ms, 44.43ms, 51.73ms, respectively. ROI, region of interest; F, fibrosis stage.

**Table 2:**
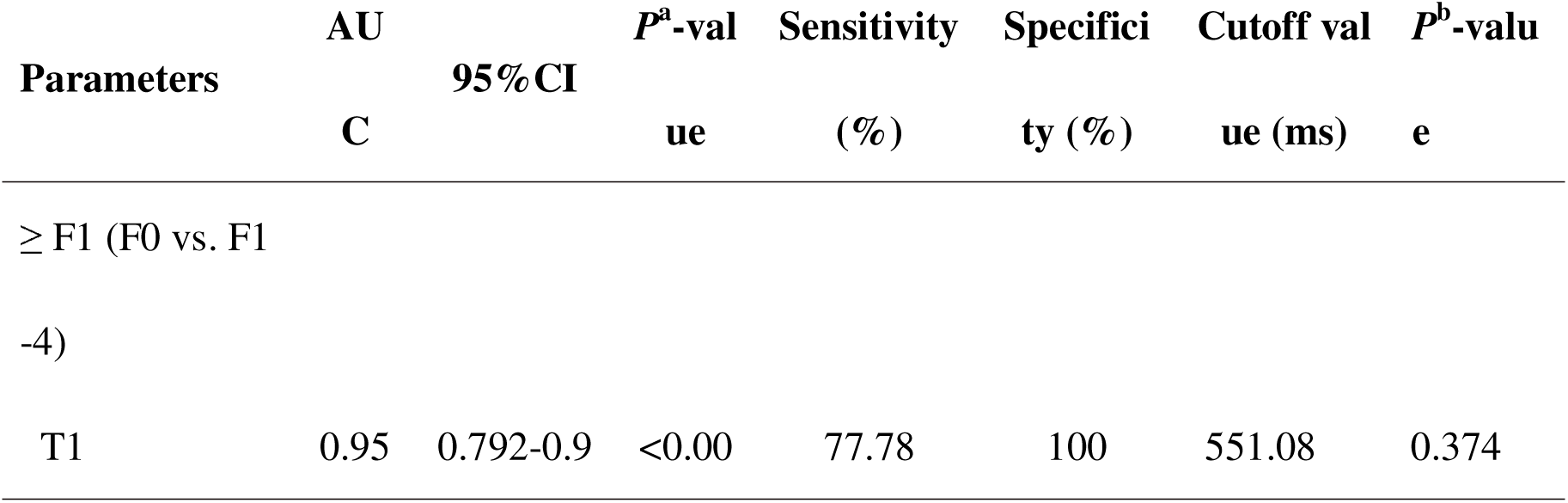
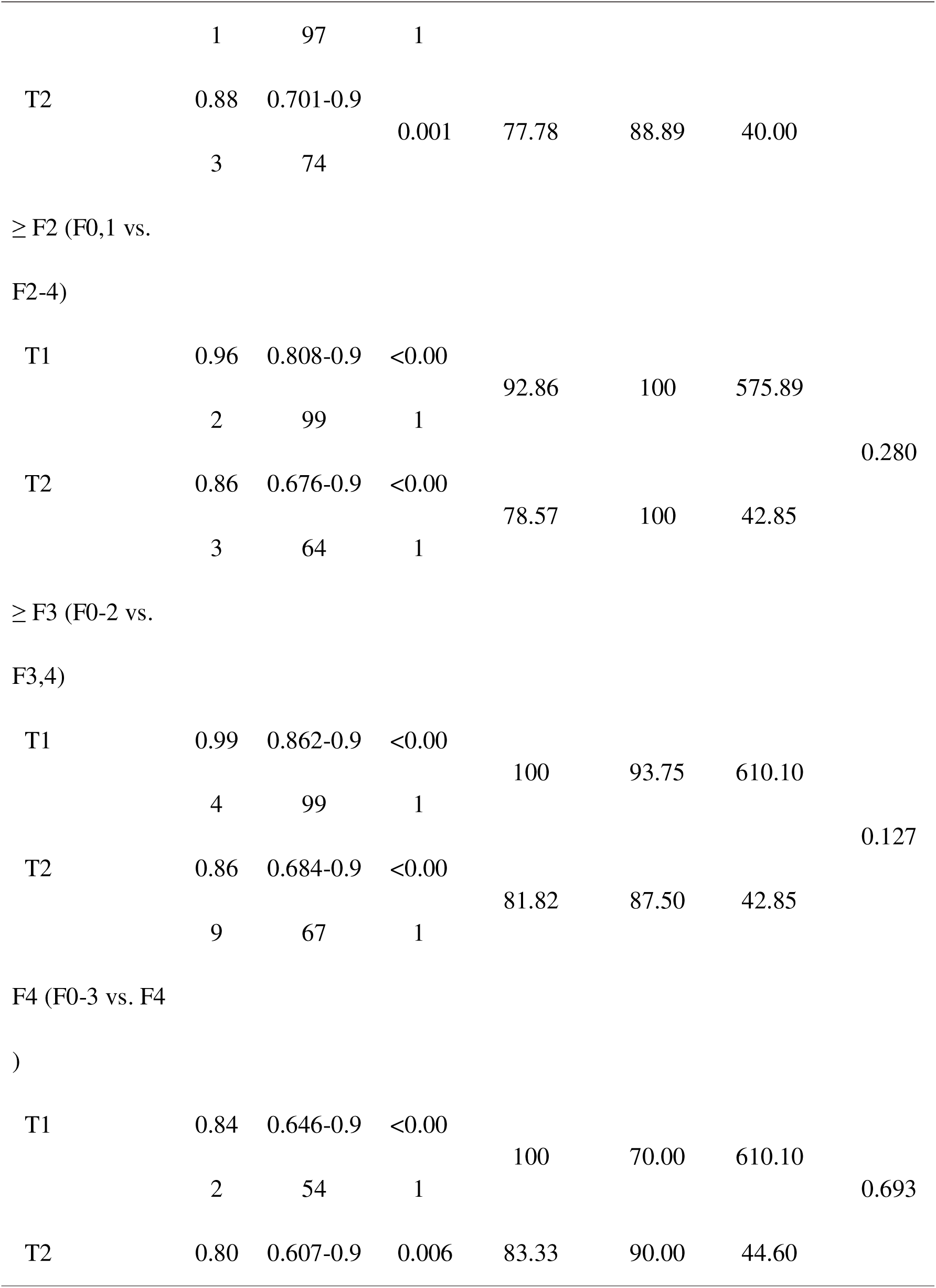
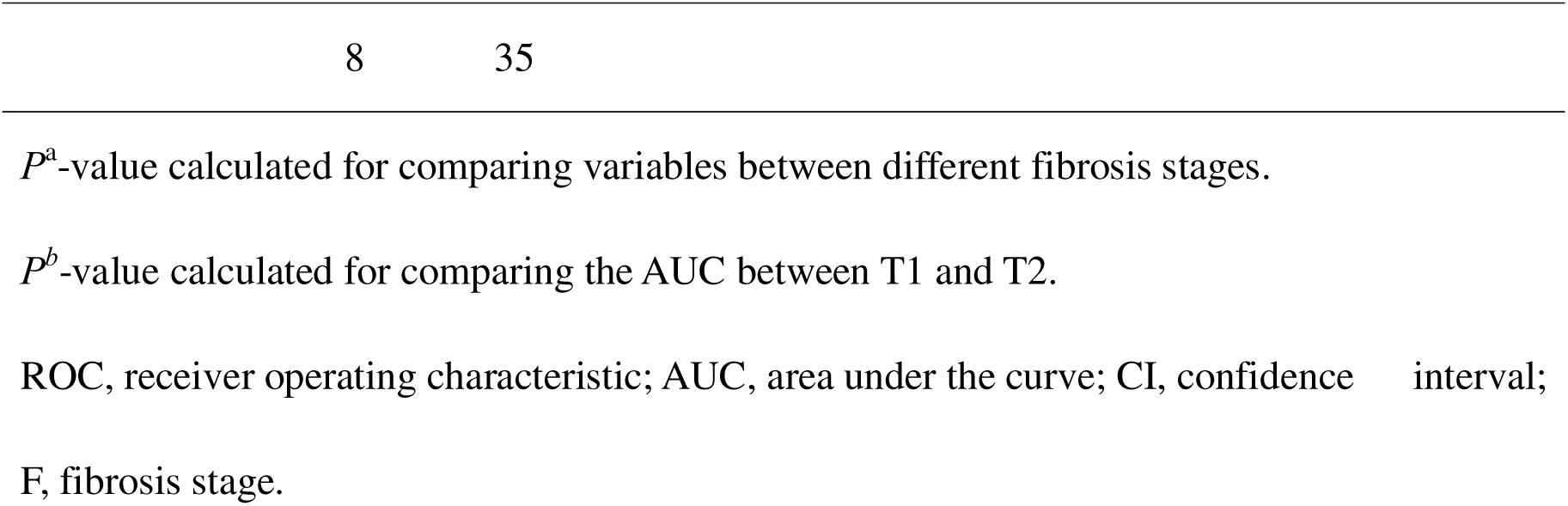
ROC curve analysis of T1 and T2 for diagnosing ≥ F1, ≥ F2, ≥ F3, ≥ F4.

| Parameters | AUC | 95%CI | *P*<sup>a</sup>-value | Sensitivity (%) | Specificity (%) | Cutoff value (ms) | *P*<sup>b</sup>-value |
| --- | --- | --- | --- | --- | --- | --- | --- |
| ≥ F1 (F0 vs. F1) | | | | | | | |
| -4) |  |  |  |  |  |  |  |
| T1 | 0.95 | 0.792-0.9 | <0.00 | 77.78 | 100 | 551.08 | 0.374 |
|  | 1 | 97 | 1 |  |  |  |  |
| T2 | 0.88 | 0.701-0.9 |  |  |  |  |  |
|  | 3 | 74 | 0.001 | 77.78 | 88.89 | 40.00 |  |
| ≥ F2 (F0,1 vs. |  |  |  |  |  |  |  |
| F2-4) |  |  |  |  |  |  |  |
| T1 | 0.96 | 0.808-0.9 | <0.00 |  |  |  |  |
|  | 2 | 99 | 1 | 92.86 | 100 | 575.89 |  |
|  |  |  |  |  |  |  | 0.280 |
| T2 | 0.86 | 0.676-0.9 | <0.00 |  |  |  |  |
|  | 3 | 64 | 1 | 78.57 | 100 | 42.85 |  |
| ≥ F3 (F0-2 vs. |  |  |  |  |  |  |  |
| F3,4) |  |  |  |  |  |  |  |
| T1 | 0.99 | 0.862-0.9 | <0.00 |  |  |  |  |
|  | 4 | 99 | 1 | 100 | 93.75 | 610.10 |  |
|  |  |  |  |  |  |  | 0.127 |
| T2 | 0.86 | 0.684-0.9 | <0.00 |  |  |  |  |
|  | 9 | 67 | 1 | 81.82 | 87.50 | 42.85 |  |
| F4 (F0-3 vs. F4 |  |  |  |  |  |  |  |
| ) |  |  |  |  |  |  |  |
| T1 | 0.84 | 0.646-0.9 | <0.00 |  |  |  |  |
|  | 2 | 54 | 1 | 100 | 70.00 | 610.10 |  |
|  |  |  |  |  |  |  | 0.693 |
| T2 | 0.80 | 0.607-0.9 | 0.006 | 83.33 | 90.00 | 44.60 |  |

$P^a$ -value calculated for comparing variables between different fibrosis stages.
$P^b$ -value calculated for comparing the AUC between T1 and T2.
ROC, receiver operating characteristic; AUC, area under the curve; CI, confidence interval;
F, fibrosis stage.

### Correlation between T1, T2 and liver fibrogenesis markers

As shown in Table 3, T1 values showed a strong correlation with HYP and α-SMA (ρ = 0.883, *P* < 0.001; ρ = 0.804, *P =* 0.002, respectively) and strong to perfect correlation with IL-1, IL-6, and TNF-α (ρ = 0.751, 0.831, 0.940, respectively; all *P* < 0.01). T2 values correlated moderately but also significantly with different liver fibrogenesis markers except for IL-1 (all *P* < 0.05).

**Table 3:** Correlation between T1, T2 and liver fibrogenesis markers.

|  | Collagen-associated protein |  |  |  | Inflammatory factors |  |  |  |  |  |
| --- | --- | --- | --- | --- | --- | --- | --- | --- | --- | --- |
| | HYP content | | α-SMA | | IL-1 | | IL-6 | | TNF-α | |
| | ρ-Val | $P$-Va | ρ-Va | $P$-Va | ρ-Va | $P$-Va | ρ-Va | $P$-Va | ρ-Va | $P$-Va |
| Parameters | ue | lue | lue | lue | lue | lue | lue | lue | lue | lue |
| T1(ms) | 0.883 | <<br>0.001 | 0.804 | 0.002 | 0.75<br>1 | 0.005 | 0.831 | 0.001 | 0.94 | <<br>0.001 |
| T2(ms) | 0.595 | 0.001 | 0.610 | 0.035 | 0.55 | 0.064 | 0.585 | 0.046 | 0.76<br>3 | 0.004 |

$P < 0.05$ indicated statistical significance.

## Discussion

This experimental animal study confirmed the usefulness of quantitative T1 and T2 mappings in the noninvasive assessment of LF at the molecular level. The strong correlations between quantitative MRI parameters and collagen-associated protein as well as inflammation factors were observed in this preclinical study.

When studying general processes LF, the CCl4-induced animal model is one of the most commonly used experimental LF models and in some ways mirrors human LF associated with toxic damage[23, 24]. Our histopathological findings demonstrated that fibrosis developed progressively from initial administration, with significant fibrosis after 2-4 weeks and severe bridging fibrosis after 5-6 weeks[24]. Corresponding to that T1 values, T2 values, and liver fibrogenesis markers increased with progressing severity of fibrosis. These findings verified the above-mentioned fundamental pathological course of LF in our CCl4-induced model, providing the possibility for us to conduct quantitative MRI techniques in the evaluation of liver fibrogenesis at different stages.

By using the CCl4-induced model in mice, we found T1 held a better diagnostic performance, sensitivity and specificity for ≥F1, ≥F2, ≥F3, F4 and correlated strongly and significantly with fibrosis stages, additionally, T1 seemed to present the best discrimination ability in the differentiation of early (F0-2) from advanced (F3-4) fibrosis. These findings were consistent with several previous studies in both animal models and human LF[13, 15]. This may likely be explained by the fact that edema and inflammation dominate early fibrosis, while inflammation and collagen deposition dominate advanced fibrosis in the CCl4-induced model[23]. In future, it needs to be verified in further clinical studies. However, using the same animal model, Chow et al. found that T1 mapping was able to distinguish normal liver from early LF, but no significant positive correlation between T1 values and fibrosis stage was observed[25]. The reason for the difference may be the different criteria used to stage fibrosis, as CCl4-induced LF is known to be associated with edema and inflammation, and there may be some overlaps between the various stages[26]. Similarly, T2 values have also reported some inconsistent results regarding the usefulness of assessing LF[11, 27–29]. In our study, T2 also presented high diagnostic performance and correlated moderately and significantly with fibrosis stages, even though not as good as the T1. This result went along with the findings of other studies evaluating T2 values in animal models and human hepatic disease, indicating that T2 may be a contributory index in assessing LF. However, a study showed a fair correlation between T2 and fibrosis stages and no significant statistical differences in T2 among fibrosis stage F0-4 in the CCl4 rat model, suggesting that T2 mapping may not be suitable for evaluating LF[13]. Several potential reasons causing the different results could be artificial bias in the ROI selection, field-strength discrepancy and experimental animal difference.

In the formation and development of LF, activated HSCs express α-SMA and are the key factors in fibrogenesis[2]. HYP is almost unique to the connective tissue collagen which is the major protein deposited in ECM[30]. Our study revealed that both T1 and T2 significantly correlated with HYP and α-SMA, which were generally consistent with Luetkens et al.[11]. However, our results showed T1 had a higher correlation than T2 with α-SMA, which was inconsistent to some extent with Luetkens et al. This may be due to the different experimental animals (rat vs mice) and methods used for quantifying α-SMA (Immunohistochemical vs Western Blot) in these two studies.

Inflammation is another core factor of LF[31]. Nuclear factor-κB (NF-κB) and other signal pathways are activated in injured hepatocytes, which induce the release of pro-inflammatory cytokines such as TNF-α, IL-1 and IL-6, and leads to the inflammatory response[32]. Our results showed a strong correlation for T1 and a moderate correlation for T2 values with inflammation factors, which was opposite to a previous mice model study by Liu et al.[17]. This discordance can be explained by the different animal models used in these two studies (the partial bile duct ligation mouse model in their study vs the CCl4 model in our study), highlighting the importance of exploring the right models to assess fibrosis. Another potential reason could be that we performed Western Blot analysis on inflammatory factors in our animal model, instead of histopathological analysis on the liver sections in their study, which would avoid the sample selection bias for inflammation analysis. This combination of findings provided some support from the perspective of molecules for the fact that T1 and T2 mapping are reliable tools for noninvasively assessing LF.

There are some limitations in this study. First, a 1.5T MRI scanner used in this study was less powerful than high field-strength or specific small-animal MRI. However, it proved to be advantageous for the easier translation of results to clinical human studies[17]. Furthermore, the effect of abdominal breathing artifacts of mice could not be completely avoided, which would reduce image quality and lead to measurement bias. Second, although the use of CCl4-induced mice models in the study of LF is a powerful tool, it may not accurately reflect the complicated histopathological changes in the human liver. Third, steatosis and iron deposition, frequently seen in humans, were not corrected in our animal model. So, care must be taken when directly transferring the above-mentioned mapping techniques to real clinical practice.

In conclusion, this study demonstrated the potential of using T1 and T2 mappings in the evaluation and differentiation of LF stages in the CCl4-induced model. T1 correlated better with collagen deposition and inflammation of LF than T2.

## Acknowledgements

We are grateful to Dr. Tianyu Yang and Dr. Guodong Zhang (Department of Pharmaceutics, School of Pharmacy, Anhui Medical University, Hefei, China) for their modelling and histopathological analysis support of this study.

## Funding

This work was supported by the National Natural Science Foundation of China (82370641) and Major Research Foundation of Higher Education of Anhui Province (2023AH040370).

## Author contributions

Yujie Zhu: study conception and design, acquisition of data, analysis and interpretation of data, drafting of manuscript, critical revision; Jing Rong: study conception and design, acquisition of data; Kun Zhu and Min Shao: acquisition of data, analysis and interpretation of data; Xiujuan Yin and Tongtong Liu: acquisition of data, analysis and interpretation of data; Xiao Wang: study conception and design, acquisition of data, analysis and interpretation. of data, critical revision.

## Conflict of interest

The authors have no competing interests to declare.

## Ethical standard

All animal experiments complied with the ARRIVE guidelines and were carried out in accordance with the National Research Council’s Guide for the Care and Use of Laboratory Animals.

## Notes

### Competing Interest Statement

The authors have declared no competing interest.

